# Visualizing Antineoplastic Activity with a Whole Organism *Drosophila melanogaster* Screen

**DOI:** 10.1101/2021.09.01.458511

**Authors:** Tristan A. Sprague, Prince N. Agbedanu

## Abstract

Cancer is a disease characterized by high mitosis rates with a loss of regulation. Many antineoplastics, those drugs used to treat cancer, act by slowing or halting mitosis. We are developing a whole-organism screening protocol to identify novel antineoplastics. After exposing *Drosophila melanogaster* eggs and larva to a compound, their growth rate and population decrease if mitosis inhibition or arrest occur. We screened several compounds from the National Cancer Institute’s (NCI) Developmental Therapeutics Program (DTP). Our screen successfully identified two compounds, toyocamycin and stictic acid, previously identified as possible antineoplastics. Toyocamycin killed a fraction of the population proportional to the dose concentration resulting in full mortality at 100 and 200 μM. At low doses, toyocamycin also slowed larval development by a mean of one day. RNAseq showed that no genes were differentially expressed in mature flies after toyocamycin exposure was halted. Stictic acid delayed larval growth by an equal or greater margin compared to toyocamycin. These results demonstrate that decreases in *Drosophila* growth or population can predict a compound’s antineoplastic activity and toxicity.

## Introduction

A plethora of screens for drug discovery and design exist, especially for novel antineoplastics. *In silico* and most *in vitro* screens require prior knowledge of a potential drug target, typically a protein or genetic sequence. Many diseases, including some cancers, are poorly characterized at the molecular level, rendering *in silico* methods and most *in vitro* methods useless. *In vivo* screens look for phenotype production or rescue making them ideal for drug discovery for less-understood diseases. Additionally, *in vitro* assays cannot recreate the cellular microenvironments present in an organism while whole-organism screens more accurately portray a compound’s biological activity and toxicity (Strange, 2016). For example, Markstein et al (2014) discovered that class II chemotherapeutics may fuel tumor reoccurrence, a potential side effect only uncovered because both cancerous and wild-type intestinal stem cells were present in the *Drosophila* model. While *Drosophila* cancer models have been used (Levine & Cagan, 2016; Rozehnal et al, 2016; Markstein et al, 2014, Rudrapatna et al, 2011) to discover successful antineoplastics in the past, we have developed a screen that does not require generation of a cancer-related genotype. This screen is an inexpensive, widely accessible technique enabling compound characterization in physiological context.

The purpose of antineoplastics is to stop the rampant cell division characterizing cancer tumors. By testing chemical libraries in whole organisms for an ability to produce slow growth or lethality these inhibitors of cell division can be identified (Giacomotto & Segalat, 2010), although morphology can be an important indicator (Wang et al, 2009). Cell division inhibition will be most noticeable where swift growth is ordinarily expected. Thus, screens using organisms in rapid stages of growth will detect such compounds with greater sensitivity than if organisms in slower stages of growth (i.e., adult *Drosophila*) were used. Our assay exposes *Drosophila* larva to potential drug candidates directly after emergence from the egg stage.

Many antineoplastics work by altering gene expression, either indirectly (e.g., through transcription factors phosphorylation) or by directly interrupting nucleic acid synthesis. For example, COX inhibitors are known to alter expression of many proteins in addition to their anti-inflammatory effects (Wang et al, 2011). 5-fluoroacil, a thymine antimetabolite, has been shown to suppress miR-200b expression in tumors (Rossi et al, 2007). Such changes in gene expression should be quantifiable by differential expression (DE) analysis of RNA sequencing data. Here, we investigate possible gene expression effects of the top hit found by our screen.

We were especially interested in gene expression effects long after drug exposure was terminated. The altered gene expression inherent to many antineoplastics could be detrimental to normal physiology if it persists long after treatment has ended. For example, Kurihara et al (2002) discovered that toyocamycin strongly increases p16/INK4a expression, thereby inhibiting cell cycle progression. While this contributes to toyocamycin’s antitumor capability, cell senescence can also delay growth. Here, we demonstrate that low-concentration toyocamycin does delay growth, but the effect is short-lived once exposure terminates.

Our protocol correctly identified toyocamycin, a nucleoside isolated from *Streptomyces* (Ohkuma, 1961), and stictic acid, a β-orcinol depsidone isolated from the lichen *Usnea articulata* (Lohézic-Le Dévéhat et al, 2007), as positive hits out of a panel provided by the NCI Developmental Therapeutics Program (DTP). Toyocamycin is a pyrrolo[2,3-d]pyrimidine (7-deazapurine), which are known to have antitumor and antiviral properties (Perlikova & Hocek, 2017) in addition to its antibiotic properties (Ohkuma, 1961). Nishioka et al (1990) recorded phosphatidyl kinase inhibition by toyocamycin. Through these mechanisms, toyocamycin perturbs nucleic acid synthesis and translation.

Stictic acid has been shown to inhibit growth in MCF-7 breast cancer and HT29 colon cancer cell lines compared to MRC-5 normal cells (Pejin, 2017). Wassman et al (2012) discovered stictic acid reactivates mutant p53. Stictic acid derivatives also have antioxidant activity against ROS (Lohézic-Le Dévéhat et al, 2007). We report similar growth inhibition in *Drosophila* in both stictic acid and toyocamycin treatments.

## Results

Toyocamycin results in death or delayed growth of *Drosophila* from the larval to adult stages. 200 and 100 micromolar toyocamycin resulted in 100% larval mortality (P=0.1233, fig. 1). Toyocamycin at 25 micromolar showed a slight reduction in time to pupation and time to emergence. Additionally, a non-significant decrease in mean population (P>.05) was observed (fig. 1). Toyocamycin and stictic groups showed a mean delay of 1 day in pupation (fig. 2–5); some stictic acid trials showed a delay of 3 days before pupation (individual data not shown). There was no significant difference in the mean population gender proportion between the treatment and control groups (P = 1) in either stictic acid or toyocamycin groups.

**Fig 1.**
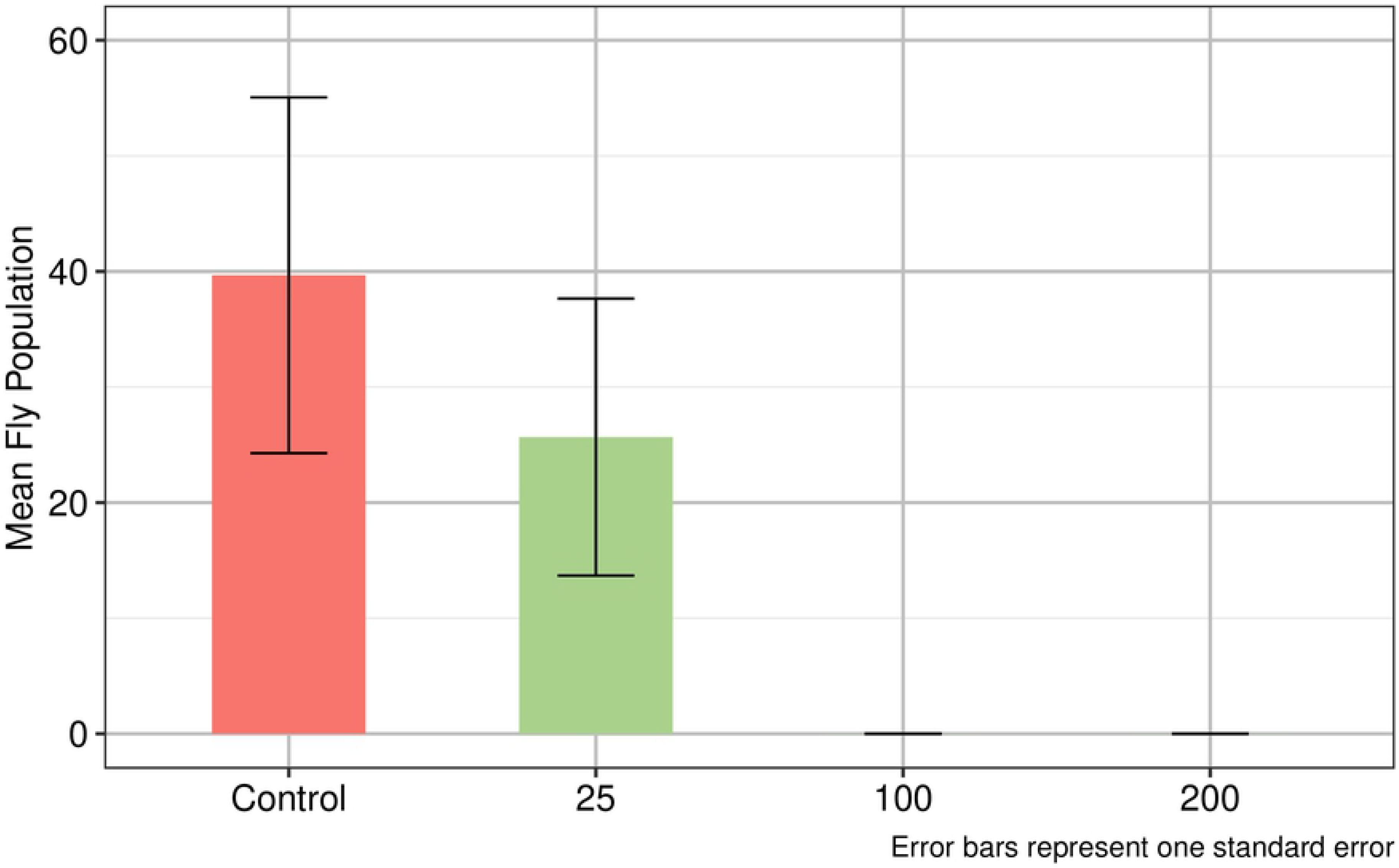
Number of viable flies decreases as toyocamycin concentration increases. Flies exposed to > 100 μM toyocamycin at the egg and larval stage saw complete mortality. Flies exposed to 25 μM toyocamycin saw a nonsignificant level of mortality.

**Fig 2.**
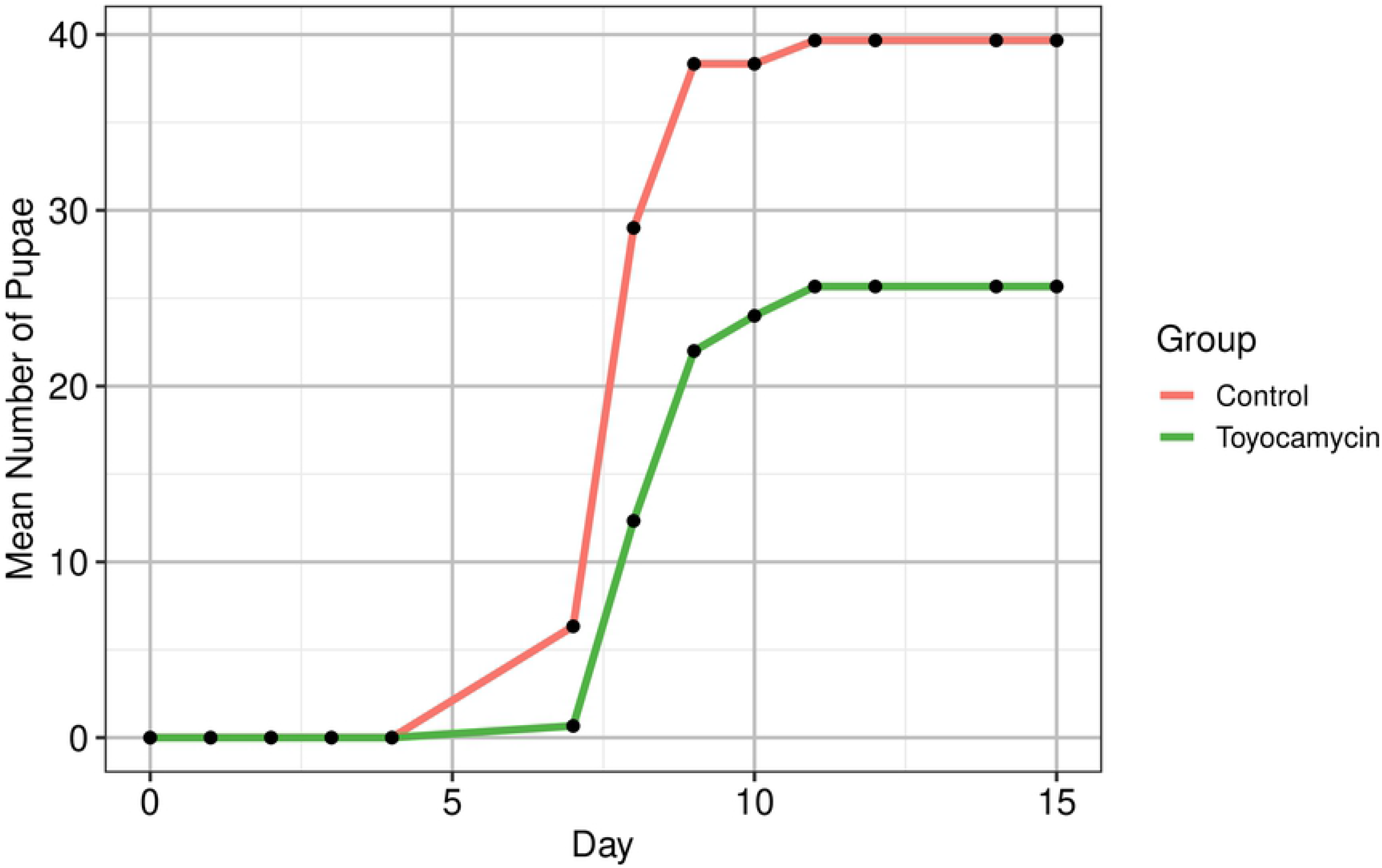
Toyocamycin, 25 μM, delays pupation of larvae. The cumulative number of pupae visible in each vial, averaged over treatments. Toyocamycin slows the appearance of pupae.

**Fig 3.**
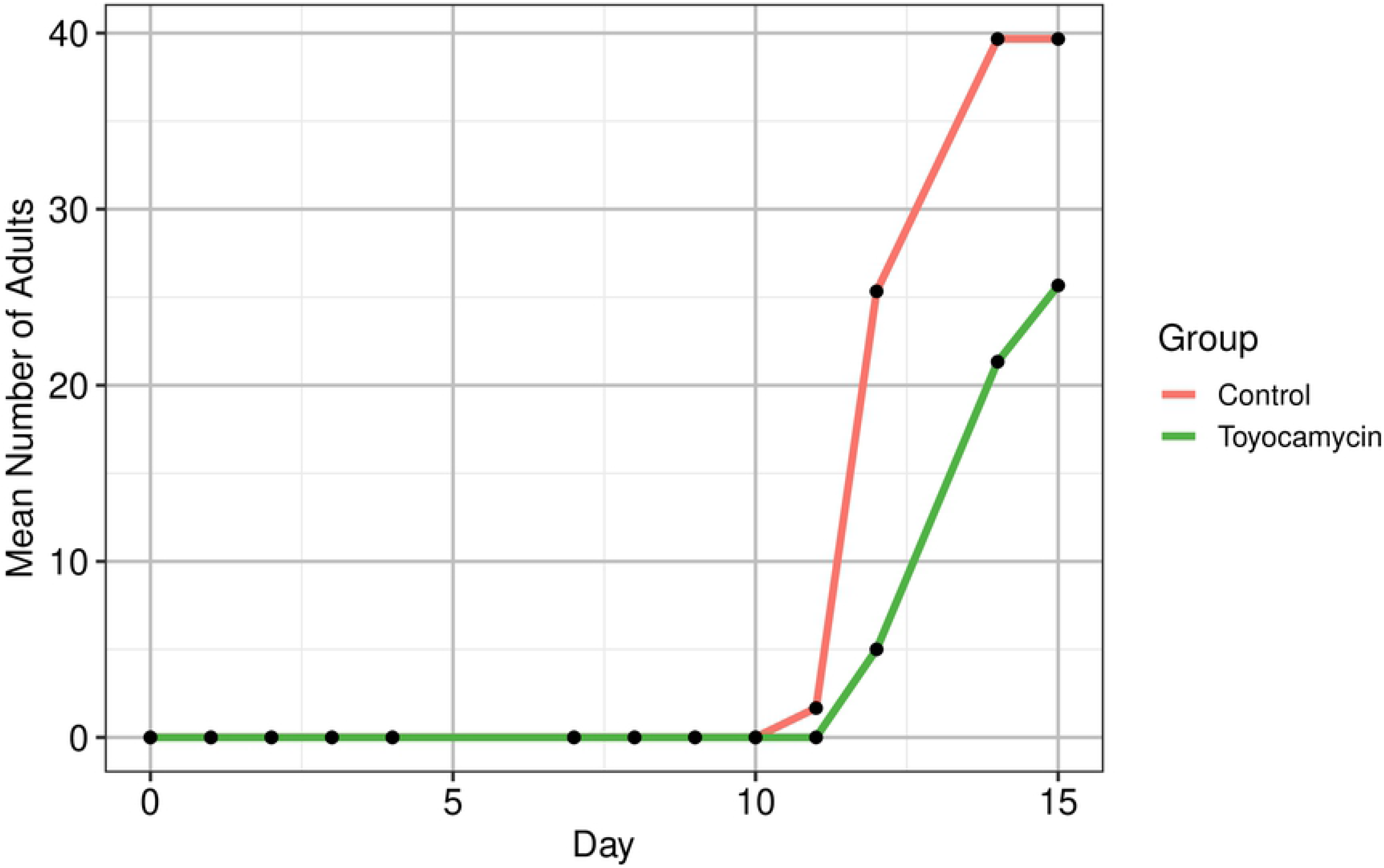
Toyocamycin, 25 μM, delays adult emergence. The cumulative number of adults visible in each vial, averaged over treatments. Toyocamycin slows the emergence of adults.

**Fig 4.**
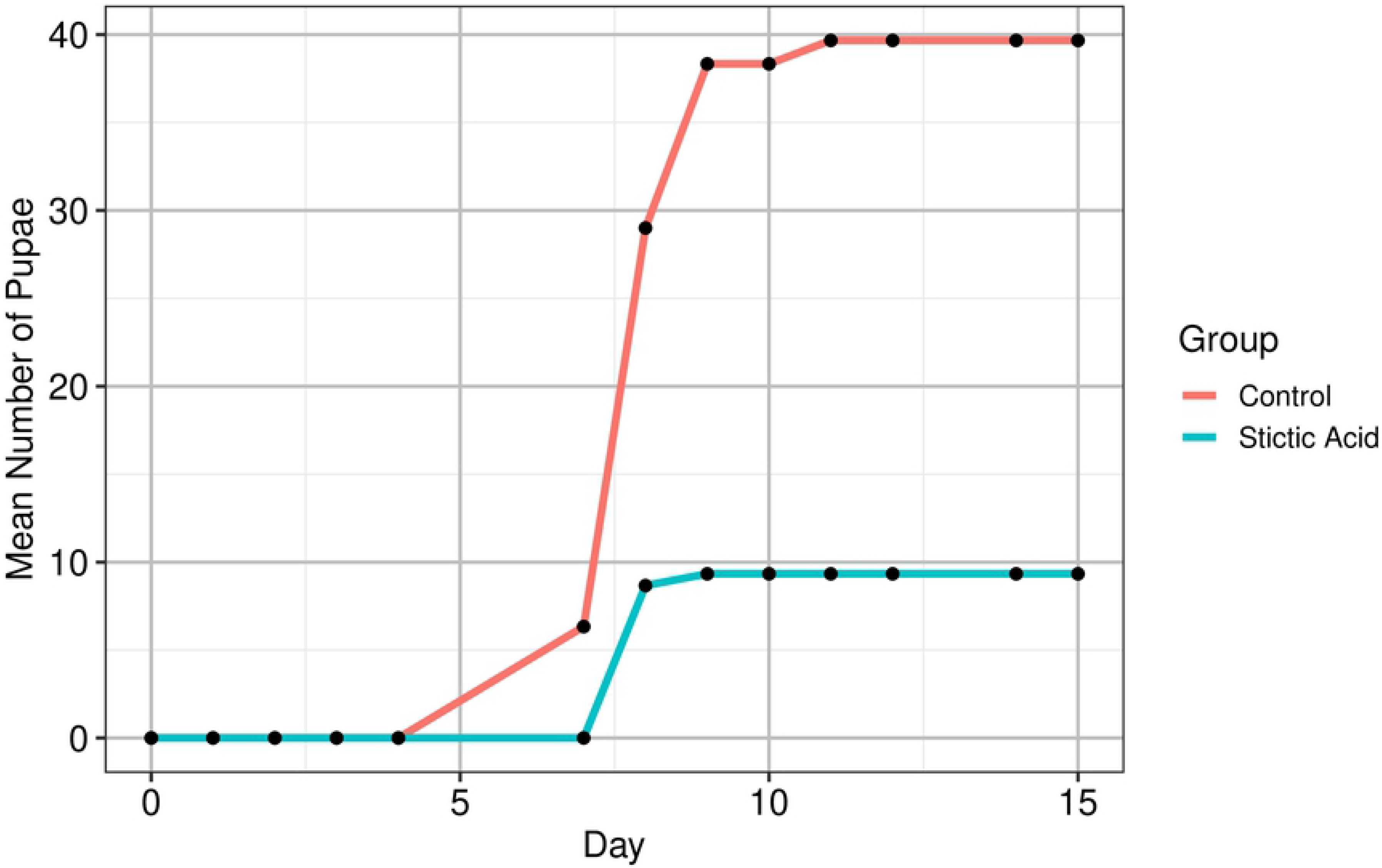
Stictic acid, 200 μM, delays pupation of larvae. The cumulative number of pupae visible in each vial, averaged over treatments. Stictic acid slows the appearance of pupae.

**Fig 5.**
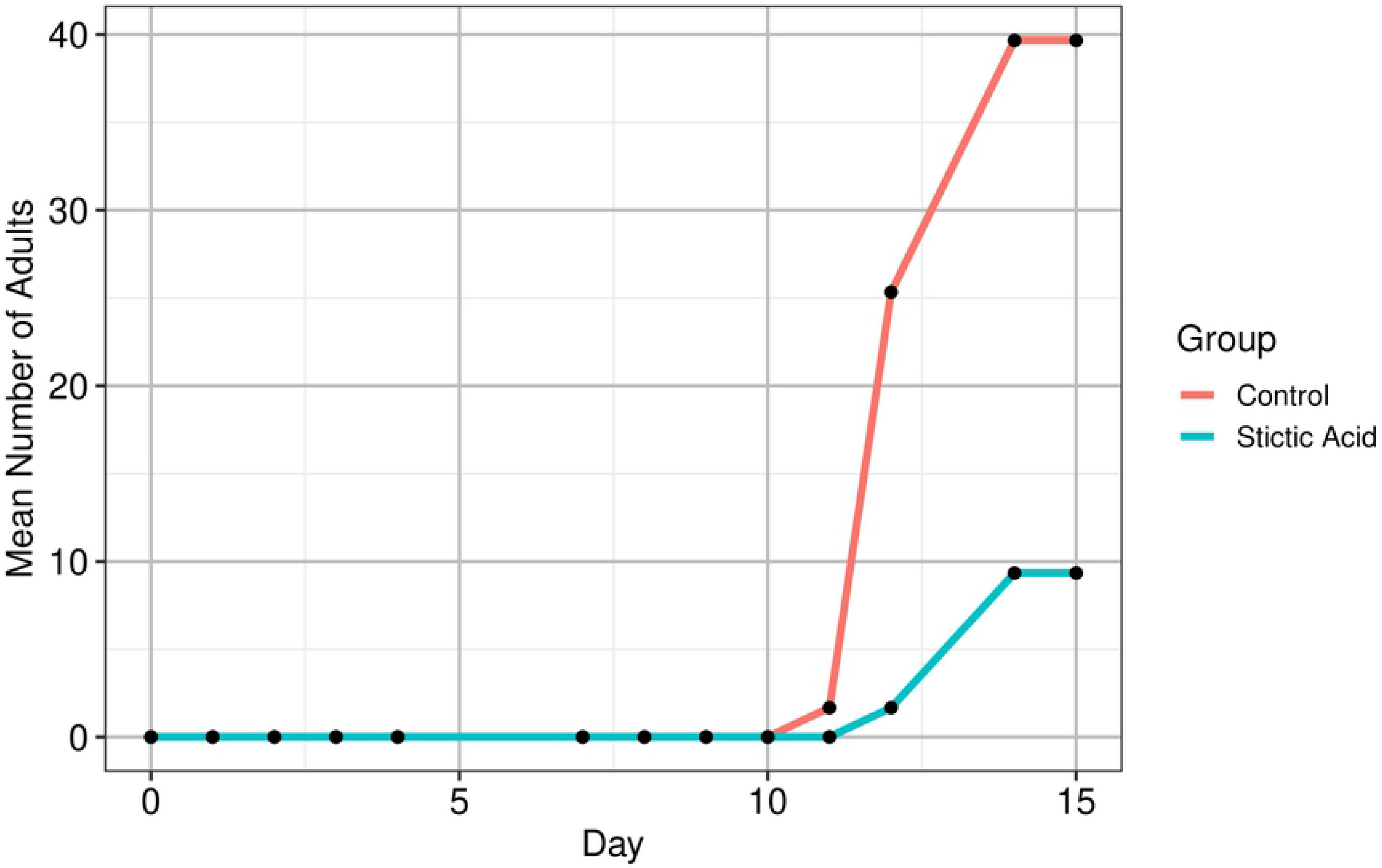
Stictic acid, 200 μM, delays adult emergence. The cumulative number of adults visible in each vial, averaged over treatments. Stictic acid slows the emergence of adults.

Preliminary RNAseq data showed 5 genes were differentially expressed significantly; CG14042, CG14933, Fbp2, and lmgA were upregulated (P=0.0474, P=0.0181, P=0.0012, P=0.0017 respectively) and snRNA:U1:95Ca was downregulated (P=0.023) (fig. 6). lmgA had the highest log2-fold change as calculated by DESeq2, 8.065 (fig. 6).

**Fig 6.**
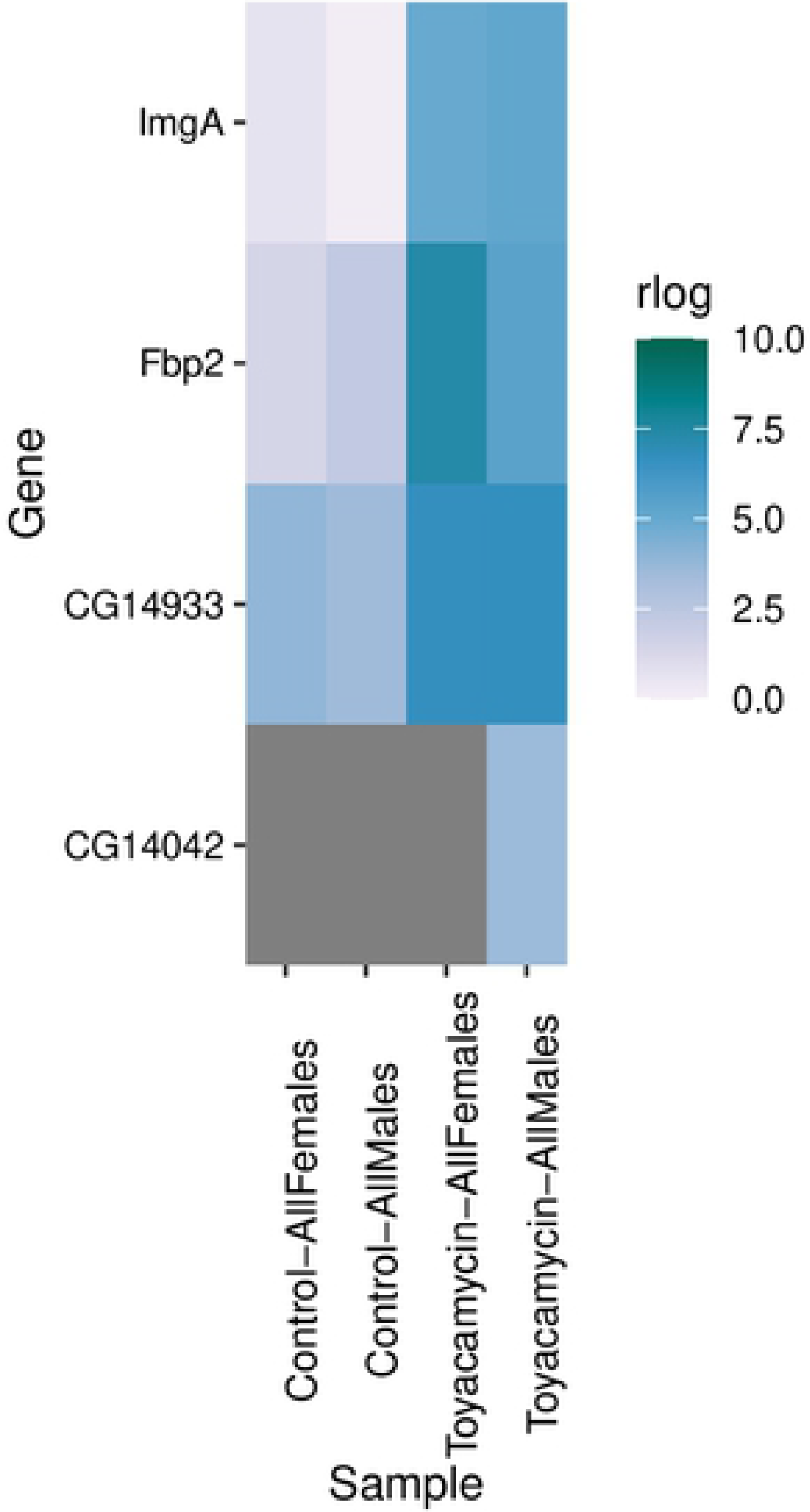
DE genes from preliminary RNAseq of toyocamycin-treated flies. RNAseq performed on male and female from both control and toyocamycin groups, (n = 3 flies per sex, with RNA combined before sequencing).

Competitive ELISA demonstrated that lmgA was not significantly DE in the toyocamycin group compared to control (fig. 7, N = 6 flies per group, P = 0.1075). Specimens used in ELISA assay were from trials independent of those used for preliminary RNAseq. Thus, another set of toyocamycin and control trials was run with a greater number of specimens sent for RNAseq.

**Fig 7.**
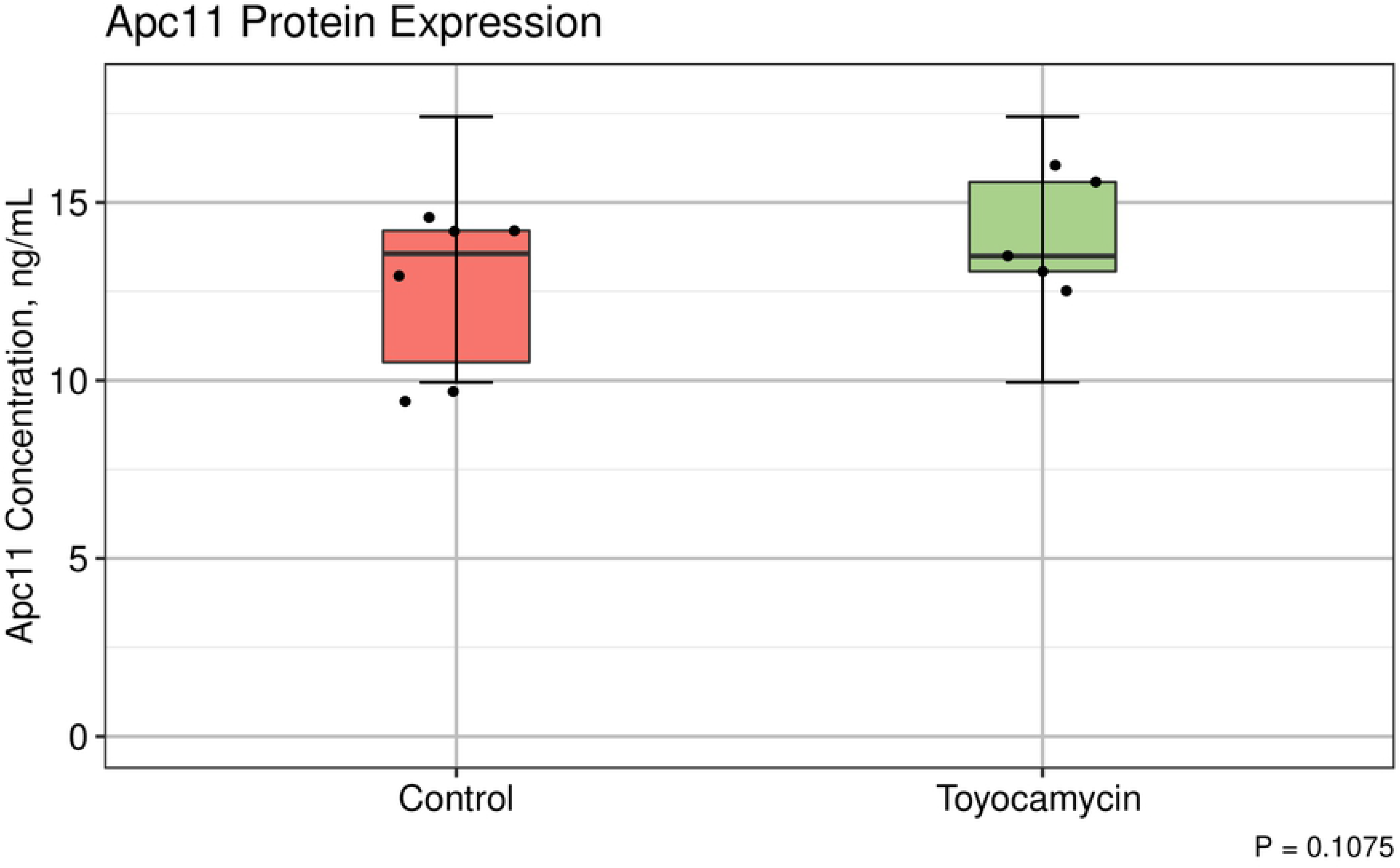
ELISA of Apc11 (lmgA) shows that lmgA was not upregulated in toyocamycin-treated flies. Competitive ELISA of Apc11 encoded by lmgA. Apc11 was not upregulated as preliminary RNAseq suggests.

Follow-up RNAseq with greater replicates (n = 6 per group) indicated no significant DE in toyocamycin treated flies at maturity. However, when flies in the toyocamycin group were compared to control flies of the same sex, 4 genes were significantly DE in males and none in females (fig. 8). However, there are only three replicates per group when sex is considered, which does not meet Schurch et al’s (2016) suggested number of biological replicates for RNAseq studies. More extensive study with greater read depth is needed to determine if sex-related differences in gene expression occur after toyocamycin exposure. It is likely that the effect we observed is due to small sampling size and interindividual variability.

**Fig 8.**
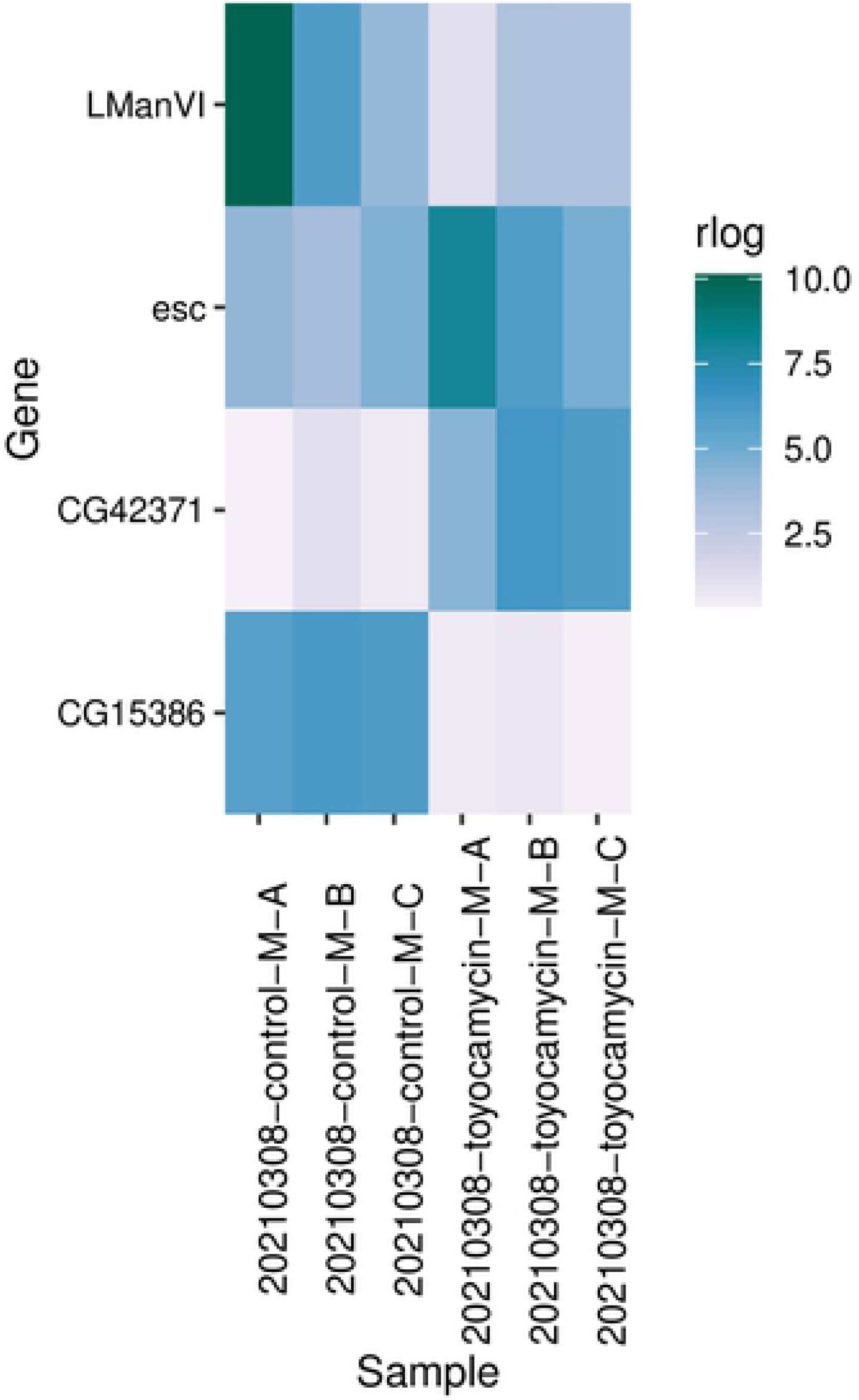
DE genes from more extensive RNAseq of male-only toyocamycin treated flies. Further RNAseq, showing DE between males in control and toyocamycin treatments (n = 3 flies per biological replicates, number of replicates = 3). When sex is ignored, no DE genes were found.

## Discussion and Conclusions

Our results show toyocamycin and stictic acid are potent inhibitors of actively growing tissues such as tumors due to the distinct response in *Drosophila*. Toyocamycin clearly inhibits cell division in *Drosophila melanogaster*, as demonstrated by the increased time by larva to reach milestones like pupation and emergence. The inhibitory effect quickly becomes toxic as concentration increases beyond 25 micromolar. Moreover, toyocamycin appears to effect male and female organisms equally. Stictic acid delayed pupation by the same margin as toyocamycin (fig. 4–5). This agrees with Pejin et al’s (2017) observations in cell lines, where clear growth inhibition was noted. These results show that our screening protocol can identify cell division inhibitors by observable effects on a developing *Drosophila* population.

While a preliminary RNAseq experiment suggested some differences in expression in mature flies after toyocamycin exposure, further analysis showed no difference. In turn, this suggest that toyocamycin ceases to affect gene expression relatively quickly when exposure stops. Furthermore, pupation did not lag far behind controls in the toyocamycin exposed flies. These flies typically pupated only one day after those in the control, while as flies in some stictic acid trials saw a delay of 3 days. Our results suggest toyocamycin’s affect fades rapidly after discontinuation. For patients undergoing chemotherapy, new drugs without long-lasting side effects once treatment is stopped are desirable. Further study of toyocamycin analogues at various time points after exposure is needed. Additionally, RNAseq with larger read depth may reveal more DE genes.

Our screen successfully identified two known antineoplastics. Additionally, new gene expression changes related to toyocamycin exposure were recorded. This was achieved without the time-consuming and expensive genetic manipulation needed to produce a cancerous genotype or patient avatar in the flies as in Markstein et al (2014) or Levine & Cagan’s (2016) work. Our screen represents a low-cost platform for studying antineoplastics *in vivo*. This technique may be used as an initial screen for suspected antineoplastics, or to elucidate the mechanism of known antineoplastics in a whole organism. Furthermore, our gene expression experiments imply that toyocamycin-based antineoplastics may have little long-term toxicity.

## Materials and Methods

### Drosophila Genetics and Maintenance; Chemical Library

Wild-type strain 25210 DGRP 859 flies were used from the Bloomington Drosophila Stock Center (Illinois, USA). Stocks were maintained on commercial media (Formula 424, Carolina Biological, North Carolina, USA) in clear plastic vials 1 1/4” diam × 4” height. Flies were moved to new vials every 30 days.

All compounds were sourced through the Drug Synthesis and Chemistry Branch, Developmental Therapeutics Program, Division of Cancer Treatment and Diagnosis, National Cancer Institute (Maryland, USA). Compounds were stored in crystalline form at −70°C until just prior to use.

### In Vivo Drug Screening

15 adult male and 15 adult females were added to each vial and treatment groups consisted of three vials each. All compounds were dissolved in DMSO (HPLC grade; ThermoFisher, Massachusetts, USA) and diluted to 200 μM, unless otherwise specified. In each vial, 15mL of dry media was reconstituted with 13 mL of 200 μM drug solution. Control vials were prepared with 15 mL of dry media and 13 mL of vehicle control. Adult flies were introduced to media for 24 hrs and then removed. Vials were maintained at 21°C with 12 hours lighting per day. 200 μM drug solutions were stored at −20°C.

An additional 2 mL of drug solution or DI water was added to each vial at 24, 48, and 72 hrs from addition of adults. Each vial was observed and the number of pupae and adult offspring recorded daily. Flies were examined for abnormal phenotypes. After emergence of all pupae, the adult vial populations were sexed and counted.

### RNAseq and Differential Expression Analysis

Specimens were randomly selected from each vial population and frozen in 1 mL RNAlater (ThermoFisher). Preliminary RNAseq specimens contained one fly per 1 ml RNAlater; triplicates were combined due to low RNA yield. Followup RNAseq specimens were allocated three flies with 1 mL RNAlater per replicate to ensure adequate yield. Specimens were stored at −70°C and shipped overnight on dry ice.

RNAseq was performed by Omega Bioservices (Georgia, USA) using Illumina sequencers at a 10M read sequencing depth. DESeq2 was utilized for differential expression analyses. DESeq2 analysis was performed using the Illumina BaseSpace platform by Omega Bioservices staff.

### Apc11 protein ELISA

To verify upregulation of lmgA past the transcription stage, ELISA of the protein Apc11 was undertaken. Samples were prepared by grinding single whole flies (n=6 with equal numbers of each sex flies per treatment.) into 500 μL PBS each using glass tissue grinders in a wet ice bath. Each sample was subjected to three freeze-thaw cycles in liquid N_2_ at room temperature. All samples were centrifugated at 5000 rpm for 15 minutes. Samples were assayed in triplicate technical replications on a single competitive human ANAPC11 ELISA plate (MyBioSource, California, USA) to avoid interassay variability. Six standards ranging from 100 ng/mL to 0 ng/mL were run in triplicate. 5-parameter logistic regression calibration curves were generated using the nplr package (Commo & Bot, 2016) in R.

### Statistical Analysis

All statistical analysis and data visualization was done using R v4.0.3 (2020) using packages ggplot2 (Wickham, 2016), dplyr (Wickham et al, 2020) and nplr, except for DESeq2 analyses. Unless otherwise noted, unpaired two sample t-tests were used to calculate p-values. Confidence levels were set at 0.95.

## Acknowledgements

We extend our deep gratitude to Dr. Chunyang Li and the other staff at Omega Bioservices for their expertise in RNAseq, useful discussion, and collaboration. We are indebted to Dr. Mostafa Zamanian of the University of Wisconsin-Madison for timely guidance on RNAseq and analysis. Stocks obtained from the Bloomington Drosophila Stock Center (NIH P40OD018537) were used in this study. Also, we thank Sarah Bottorff of Carolina Biological for suggestions on controlling *Drosophila* bacterial infections.

We wish to thank the Friends University Division of Science, Technology, Engineering, and Math, especially Dr. Nora Strasser (Chair), Amy Morgan (Admin), and Celia Milam (*Drosophila* stockkeeper). Additionally, we are grateful to Ramon Emmart, Jessica Boone, Erin E. McCoskey (PA-S), and Abbey L. Fischer (CMA) for their scrutiny of our grant proposal.

Most of all, we want to thank the Kansas Academy of Science, their Undergraduate Scholarship Committee, and Dr. Erin Morris for providing us with partial funding through the Student Research Grant Program.

